# Deep Learning based multi-omics integration robustly predicts survival in liver cancer

**DOI:** 10.1101/114892

**Authors:** Kumardeep Chaudhary, Olivier B. Poirion, Liangqun Lu, Lana X. Garmire

## Abstract

Identifying robust survival subgroups of hepatocellular carcinoma (HCC) will significantly improve patient care. Currently, endeavor of integrating multi-omics data to explicitly predict HCC survival from multiple patient cohorts is lacking. To fill in this gap, we present a deep learning (DL) based model on HCC that robustly differentiates survival subpopulations of patients in six cohorts. We build the DL based, survival-sensitive model on 360 HCC patients’ data using RNA-seq, miRNA-seq and methylation data from TCGA, which predicts prognosis as good as an alternative model where genomics and clinical data are both considered. This DL based model provides two optimal subgroups of patients with significant survival differences (P=7.13e-6) and good model fitness (C-index=0.68). More aggressive subtype is associated with frequent *TP53* inactivation mutations, higher expression of stemness markers (*KRT19*, *EPCAM*) and tumor marker *BIRC5*, and activated Wnt and Akt signaling pathways. We validated this multi-omics model on five external datasets of various omics types: LIRI-JP cohort (n=230, C-index=0.75), NCI cohort (n=221, C-index=0.67), Chinese cohort (n=166, C-index=0.69), E-TABM-36 cohort (n=40, C-index=0.77), and Hawaiian cohort (n=27, C-index=0.82). This is the first study to employ deep learning to identify multi-omics features linked to the differential survival of HCC patients. Given its robustness over multiple cohorts, we expect this workflow to be useful at predicting HCC prognosis prediction.

## Introduction

Liver cancer is the 2^nd^ leading cancer responsible for the mortality in men, worldwide (1). In USA, more than 40,000 people are estimated to be diagnosed with liver cancer in 2017, according to the American Cancer Society (2). It is one of the few cancer types with increase in both incidence and mortality rates, by ~3 % per year in the US (3). Hepatocellular carcinoma (HCC) is the most prevalent type (70-90%) of liver cancer. It is aggravated by various risk factors, including HBV/HCV infection, nonalcoholic steatohepatitis (NASH), alcoholism, and smoking. The 5-year survival rate of HCC varies greatly from different populations, with an average rate of less than 32% (4-9). The high level of heterogeneity in HCC along with the complex etiological factors makes the prognosis prediction very challenging (10,11). Moreover, treatment strategies in HCC are very limited, imposing additional urgent needs for developing tools to predict patient survival (12).

To understand the HCC heterogeneity among patients, a considerable amount of work has been done to identify the HCC molecular subtypes (13-19). A variety of numbers of subtypes were identified, ranging from 2 to 6, based on various omics data types, driving hypotheses and computational methods. Besides most commonly used mRNA gene expression data, a recent study integrated copy number variation (CNV), DNA methylation, mRNA and miRNA expression to identify the 5 HCC molecular subtypes from 256 TCGA samples (20). However, most of the studies explored the molecular subtypes without relying on survival during the process of defining subtypes (21). Rather, survival information was used *post hoc* to evaluate the clinical significance of these subtypes (20). As a result, some molecular subtypes showed converging and similar survival profile, making them redundant subtypes in terms of survival differences (16). New approaches to discover survival-sensitive and multi-omics data based molecular subtypes are much needed in HCC research.

To address these issues, for the first time, we have utilized deep learning (DL) computational framework on multi-omics HCC datasets. We chose the autoencoder framework as the implementation of DL for multi-omics integration. Autoencoders aim to reconstruct the original input using combinations of nonlinear functions which can then be used as new features to represent the dataset. These algorithms have already been proved to be efficient approaches to produce features linked to clinical outcomes (22). Autoencoders were successfully applied to analyze high-dimensional gene expression data (23,24), and to integrate heterogeneous data (25,26). Notably, autoencoder transformation tends to aggregate genes sharing similar pathways (27), therefore making it appealing to interpret the biological functions. The contributions of this study to HCC field is not only manifested in its thorough and integrative computational rigor, but also unify the discordant molecular subtypes into robust subtypes that withstand the testing of various cohorts, even when they are in different omics forms.

We derived the model from 360 HCC samples in TCGA multi-omics cohort, which have mRNA expression, miRNA expression, CpG methylation and clinical information. We discovered two subtypes with significant differences in survival. These subtypes hold independent predictive values on patient survival, apart from clinical characteristics. Most importantly, the two subtypes obtained from our DL framework are successfully validated in five independent cohorts, which have miRNA or mRNA or DNA methylation dataset. Functional analysis of these two subtypes identified that gene expression signatures (*KIRT19*, *EPCAM* and *BIRC5*) and Wnt signaling pathways are highly associated with poor survival. In summary, the survival-sensitive subtypes model reported here is significant for both HCC prognosis prediction and therapeutic intervention.

## Materials and Methods

### Datasets and study design

In this study, we used a total of 6 cohorts, and the descriptions of them are detailed below. We used TCGA data in two steps: the first step is to obtain the labels of survival-risk classes, using the whole TCGA data set; another step is to train an SVM model, by splitting the samples 60/40% to training and holdout testing data (detailed in “Data partitioning and robustness assessment” subsection). We used 5 additional confirmation data sets to evaluate the prediction accuracy of the DL based prognosis model.

#### TCGA set

We obtained multi-omics HCC data from the TCGA portal (https://tcga-data.nci.nih.gov/tcga/). We used R package TCGA-assembler (v1.0.3) (28) and obtained 360 samples with RNA-seq data (UNC IlluminaHiSeq_RNASeqV2; Level 3), miRNA-seq data (BCGSC IlluminaHiSeq_miRNASeq; Level 3), DNA methylation data (JHU-USC HumanMethylation450; Level 3), and the clinical information. For the DNA methylation, we mapped CpG islands within 1500 bp ahead of transcription start sites (TSS) of genes and averaged their methylation values. In dealing with the missing values (preprocessing of data), three steps were performed as elsewhere (29). First, the biological features (e.g. genes/miRNAs) were removed if having zero value in more than 20% of patients. The samples were removed if missing across more than 20% features. Then we used *impute* function from R impute package (30), to fill out the missing values. Lastly, we removed input features with zero values across all samples.

#### Confirmation cohort 1 (LIRI-JP cohort, RNA-seq)

230 samples with RNA-seq data were obtained from ICGC portal (https://dcc.icgc.org/projects/LIRI-JP). These samples belong to Japanese population primarily infected with HBV/HCV (31). We used the normalized read count values given in the gene expression file.

#### Confirmation cohort 2 (NCI cohort, microarray gene expression)

221 samples with survival information were chosen from GSE14520 Affymetrix high-throughput GeneChip HG-U133A microarray dataset, from an earlier study of HCC patients (32). This is a Chinese population primarily associated with HBV infection. Log2 Robust Multi-array Average (RMA)-calculated signal intensity values provided by the authors were used for analysis.

#### Confirmation cohortS 3 (Chinese cohort, miRNA expression array)

166 pairs of HCC/matched noncancerous normal tissue samples were downloaded, with CapitalBio custom Human miRNA array data (GSE31384) (33). Since the data were already log2 transformed, we used unit-scale normalization.

#### Confirmation cohort 4 (E-TABM-36, gene expression microarray)

40 HCC samples were used, with survival information and transcriptional profiling from Affymetrix HG-U133A GeneChips arrays platform (16). We used the CHPSignal values for the further processing as a measure of gene expression.

#### Confirmation cohort 5 (Hawaiian cohort, DNA Methylation array)

27 samples were used, with genome-wide methylation profiling from Illumina HumanMethylation450 BeadChip platform (34). Probe to gene conversion was done the same way as for TCGA HCC methylation data.

All the available clinical information for the confirmation cohorts is listed in Supplementary Table S1. These cohorts were used to test the Support Vector machine (SVM) machine-learning models.

### Transformed features using a deep Learning framework

We used the 3 preprocessed TCGA HCC omics datasets of 360 samples as the input for the autoencoders framework. We stacked the 3 matrices that are unit-norm scaled by sample, in order to form a unique matrix as reported before (35). An autoencoder is an unsupervised feed-forward, non-recurrent neural network (36). Given an input layer taking the input *x* = (*x_1_*,…,*x_n_* of dimension *n*, the objective of an autoencoder is to reconstruct *x* by the output *x'* (*x* and *x’* have the same dimension), via transforming *x* through successive hidden layers. For a given layer *i*, we used *tanh* as activation function between input layer *x* and output layer *y*. That is:

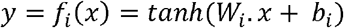

Where *x* and *y* are two vectors of size *d* and *p*, respectively and *Wi* is the weight matrix of size *d* × *p*, *b_i_* an intercept vector of size *p* and *W_i_*.*x* = ∑*W_i,j_. x_j_*, with *x_j_* the value of a single feature from *x*. For an autoencoder with *k* layers, *x´* is then given by:

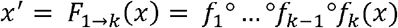

Where *f*_*k*-1_°*f_k_*(*x*) is the composed function of *f*_*k*-1_ with *f_k_*. To train an autoencoder, the objective is to find the different weight vectors *W_i_* minimizing a specific objective function. We chose *logloss* as the objective function which measures the error between the input *x* and the output *x'*:

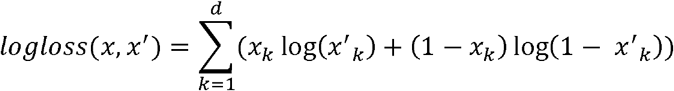

In order to control overfitting, we added an *L1* regularization penalty *α*_*w*_ on the weight vector *W_i_*, and a *L2* regularization penalty *α*_*a*_ on the nodes activities: *F*_1→*k*_(*x*). Thus the objective function above becomes:

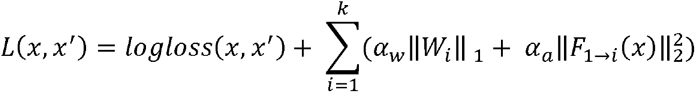

We implemented an autoencoder with three hidden layers 500, 100 and 500 nodes respectively, using python Keras library (https://github.com/fchollet/keras). We used the bottleneck layer of the autoencoder to produce new features from the omics data. The values α_*a*_ and α_*w*_ were set to 0.0001 and 0.001. Finally, to train the autoencoder we used the gradient descent algorithm with 10 epochs and 50% dropout. Here, epoch means the iteration of the learning algorithm (stochastic gradient descent) through the entire training dataset. During one epoch, the learning algorithm processes each instance of training data once.

### Transformed feature selection and K-means clustering

The autoencoder reduced the initial number of features to 100 new features obtained from the bottleneck layer. Next for each of these transformed features produced by autoencoder, we built a univariate Cox-PH model and selected features from which a significant Cox-PH model is obtained (log-rank p-value < 0.05). We then used these reduced new features to cluster the samples using the K-means clustering algorithm. We determined the optimal number of clusters with two metrics: Silhouette index (37) and Calinski-Harabasz criterion (38). We used the *scikit-learn* package for K-means implementation (39).

### Data partitioning and robustness assessment

We used a CV like procedure to partition the TCGA dataset as follows: We used a 60/40% split (training/test sets) of the TCGA data, in order to have sufficient number of test samples that generate evaluation metrics. We first randomly split the 360 samples from TCGA into 5 folds. We then used 2 of the 5 of folds as the test set, and the remaining 3 folds as the training set. With this approach, we obtained 10 new combinations (folds). For each of these 10 new folds, we constructed a model using the 60% samples (training set) and predicted the labels in test set (held-out). This data partitioning was only used to assess the robustness of the model. For each training fold, a distinct autoencoder and a classifier (see below) were built to predict the labels of the test fold. The labels of the TCGA samples are finally inferred using an autoencoder built with all the samples and these labels were used for the prediction of the confirmation datasets.

### Supervised classification

After obtaining the labels from K-means clustering, we built supervised classification model(s) using SVM algorithm. We normalized each omics layer in the training set, then selected the top N features which are most correlated with the cluster labels (obtained from K-means), based on ANOVA F-values. We set default N values as 100 for mRNAs, 50 for methylation and 50 for miRNAs.

To predict on TCGA 3 omics held-out test data, we built an SVM classifier from a combination of top 100 mRNAs, 50 methylation and 50 miRNA features, selected by ANOVA. To predict each of the other 5 confirmation cohorts used in this study, we built an SVM classifier on each omics type, using the corresponding top 100 mRNAs, 50 methylation, or 50 miRNA features selected by ANOVA, respectively. For a confirmation cohort from a specific omic layer, we first selected common features (mRNAs, CpG sites or miRNAs) between this cohort and the corresponding omic layer in the TCGA training set. Specifically, the common features between the five cohorts and TCGA training dataset are: 14634 for LIRI-JP cohort, 9311 for NCI cohort, 174 for Chinese cohort, 10550 for E-TABM-36 cohort and 19883 for Hawaiian cohort.

We then applied two scaling steps on both training set and the confirmation cohort samples. We first used a median scaling on both the training set and the new test samples, where each feature is rescaled according to its median and absolute median deviation. This approach was used to normalize samples from RNA-seq data previously (40). For mRNA and DNA methylation data, we then applied a robust scaling on the training set and confirmation samples using the means and the standard deviations of the training set (41). For miRNA confirmation data, we applied the unit scale normalization for both the miRNA training and the confirmation cohort. When predicting a single sample, an alternative rank normalization (rather than robust or unit scale normalization) can be applied to both the new sample and samples from the training sets (see more details in File S1).

We used the *scikit-learn* package to perform grid search to find the best hyperparameters of the SVM model(s) using 5-fold cross-validation (CV) and built SVM models.

### Evaluation metrics for models

The metrics used closely reflect the accuracy of survival prediction in the subgroups identified. Three sets of evaluation metrics were used.

#### Concordance index (C-index)

The C-index can be seen as the fraction of all pairs of individuals whose predicted survival times are correctly ordered (42) and is based on Harrell’s C statistics (43). A C-index score around 0.70 indicates a good model, whereas a score around 0.50 means random background. To compute the C-index, we first built a Cox-PH model using the training set (cluster labels and survival data) and predict survival using the labels of the test/confirmation set. We then calculated the C-index using function concordance.index in R *survcomp* package (44). To compute the C-index using the multiple clinical features, we built a Cox-PH using the *glmnet* package instead (45). We opted to perform penalization through ridge regression rather than the default Lasso penalization. Before building the Cox-PH model, we performed a 10-fold CV to find the best lambda.

#### Log-rank p-value of Cox-PH regression

We plotted the Kaplan-Meier survival curves of the two risk groups, and calculated the log-rank p-value of the survival difference between them. We used Cox proportional hazards (Cox-PH) model for survival analysis (46), similar to described before (47,48), using R *survival* package (49).

#### Brier score

It is another score function that measures the accuracy of probabilistic prediction (50). In survival analysis, the brier score measures the mean of the difference between the observed and the estimated survival beyond a certain time (51). The score ranges between 0 and 1 and a larger score indicates higher inaccuracy. We used the implementation of brier score from R *survcomp* package.

### Alternative approaches to the deep Learning framework

We compared the performances of the deep learning framework with two alternative approaches. In the first approach, we performed PCA analysis and used the same number (100) of principal components as those features in the bottleneck layer of Figure 1. We then identified the subset (13) of PCA features significantly associated with survival using univariate Cox-PH models, using the same procedure as the Cox-PH step in Figure 1. In the second approach, we selected top 37 features among all three omics features using single-variant Cox-PH models, based on the C-index scores. We clustered the samples using the same K-means procedure as in Figure 1.

**Figure 1:**
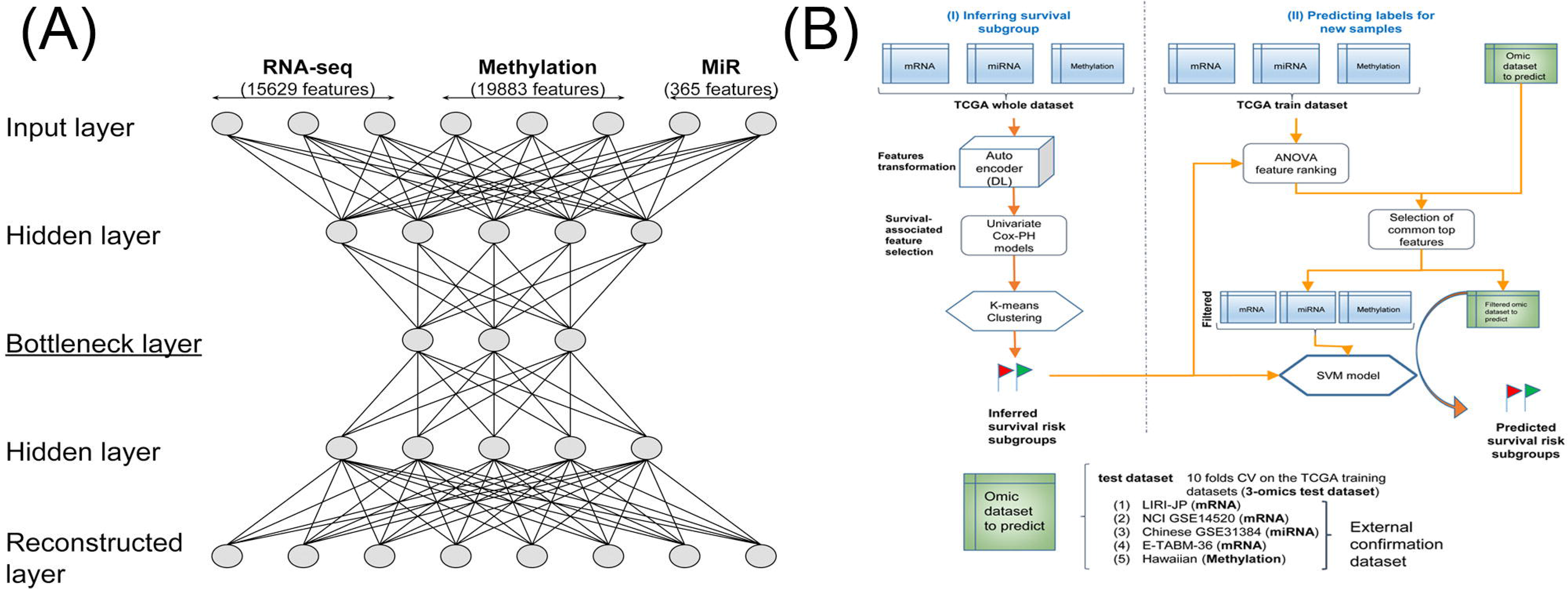
Overall workflow (A) Autoencoder architecture used to integrate 3 omics of HCC data. (B) Workflow combining deep learning and machine learning techniques to predict HCC survival subgroups. The workflow includes two steps. Step 1: inferring survival subgroups and Step 2: predicting risk labels for new samples. In step 1: mRNA, DNA methylation and miRNA features from TCGA HCC cohort are stacked up as input features for autoencoder, a deep learning method; then each of the new, transformed features in the bottle neck layer of autoencoder is then subject to single variate Cox-PH models, to select the features associated with survival; then K-mean clustering is applied to samples represented by these features, to identify survival-risk groups. In step 2, mRNA, methylation and miRNA input features are ranked by ANOVA test F-values, those features that are in common with the predicting dataset are selected, then top features are used to build SVM model(s) to predict the survival risk labels of new datasets.

### Functional analysis

A number of functional analyses were performed to understand the characteristics of 2 survival risk subtypes of TCGA HCC samples.

#### TP53 mutation analysis

We analyzed the somatic mutation frequency distributions in the survival subtypes for the *TP53* gene, among TCGA and LIRI-JP cohorts. TCGA and LIRI-JP cohorts have exome sequencing and whole genome sequencing data for 186 and 230 samples with survival data, respectively. We performed Fisher’s test on *TP53* mutation between two survival risk groups.

#### Clinical covariate analysis

We tested the associations of our identified subtypes with other clinical factors, including gender, race, grade, stage and risk factors, using Fisher’s exact tests. To test if the two survival risk subtypes have prognostic values in addition to clinical characteristics, we built a combined Cox-PH model with survival risk classification and clinical data, and compared it to the one with only clinical data (stage, grade, race, gender, age and risk factor).

#### Differential Expression

In order to identify the differential expressed genes between the two survival risk subtypes, we performed the differential expression analysis for the mRNA, miRNA expression and methylation genes. We used DESeq2 package (52) to identify the differential gene and miRNA expression between the 2 subtypes (false discovery rate, or FDR <0.05). Additionally, we used log2 fold change greater than 1 as filtering for mRNA/miRNA. For methylation data, we transformed the beta values into M values as elsewhere (53,54) using the *lumi* package in R (55). We fit the linear model for each gene using *lmFit* function followed by empirical Bayes method, using *limma* package in R (56). It uses moderate t-tests to determine significant difference in methylation for each gene between S1 and S2 subtypes (Benjamin-Hochberg corrected P<0.05). Additionally, we used averaged M value differences greater than 1 as filtering. We used volcano plot to show the differentially methylated genes in two subtypes.

#### Enriched pathway analysis

We used upregulated and downregulated genes for the KEGG pathway analysis, using the functional annotation tool from the online DAVID interface (57,58). We used the modified Fisher’s Exact Test p-value (EASE score provided by DAVID) threshold of 0.10 to consider a pathway significant. We plot the gene-pathway network using Gephi (59).

## Results

### Two differential survival subtypes are identified in TCGA multi-omics HCC data

From the TCGA HCC project, we obtained 360 tumor samples that had coupled RNA-seq, miRNA-seq and DNA methylation data. For these 360 samples, we preprocessed the data as described in the ‘Materials and Methods’ section, and obtained 15,629 genes from RNA-seq, 365 miRNAs from miRNA-seq, and 19,883 genes from DNA methylation data as input features. These three types of omics features were stacked together using autoencoder, a deep learning framework (36). The architecture of autoencoder is shown in Figure 1A. We used the activity of the 100 nodes from the bottleneck hidden layer as new features. We then conducted univariate Cox-PH regression on each of the 100 features, and identified 37 features significantly (log-rank p-value <0.05) associated with survival. These 37 features were subjective to K-means clustering, with cluster number K ranging from 2 to 6. Using silhouette index and the Calinski-Harabasz criterion, we found that K=2 was the optimum with the best scores for both metrics (Figure S1A). Further, the survival analysis on the full TCGA HCC data shows that the survivals in the two sub-clusters are drastically different (log-rank p-value =7.13e-6, Figure 2A). Moreover, K=2 to 6 yielded KM survival curves that essentially represent 2 significantly different survival groups (Figure S1B). Thus, we determined that K=2 was the classification labels for the subsequent supervised machine learning processes.

**Figure 2:**
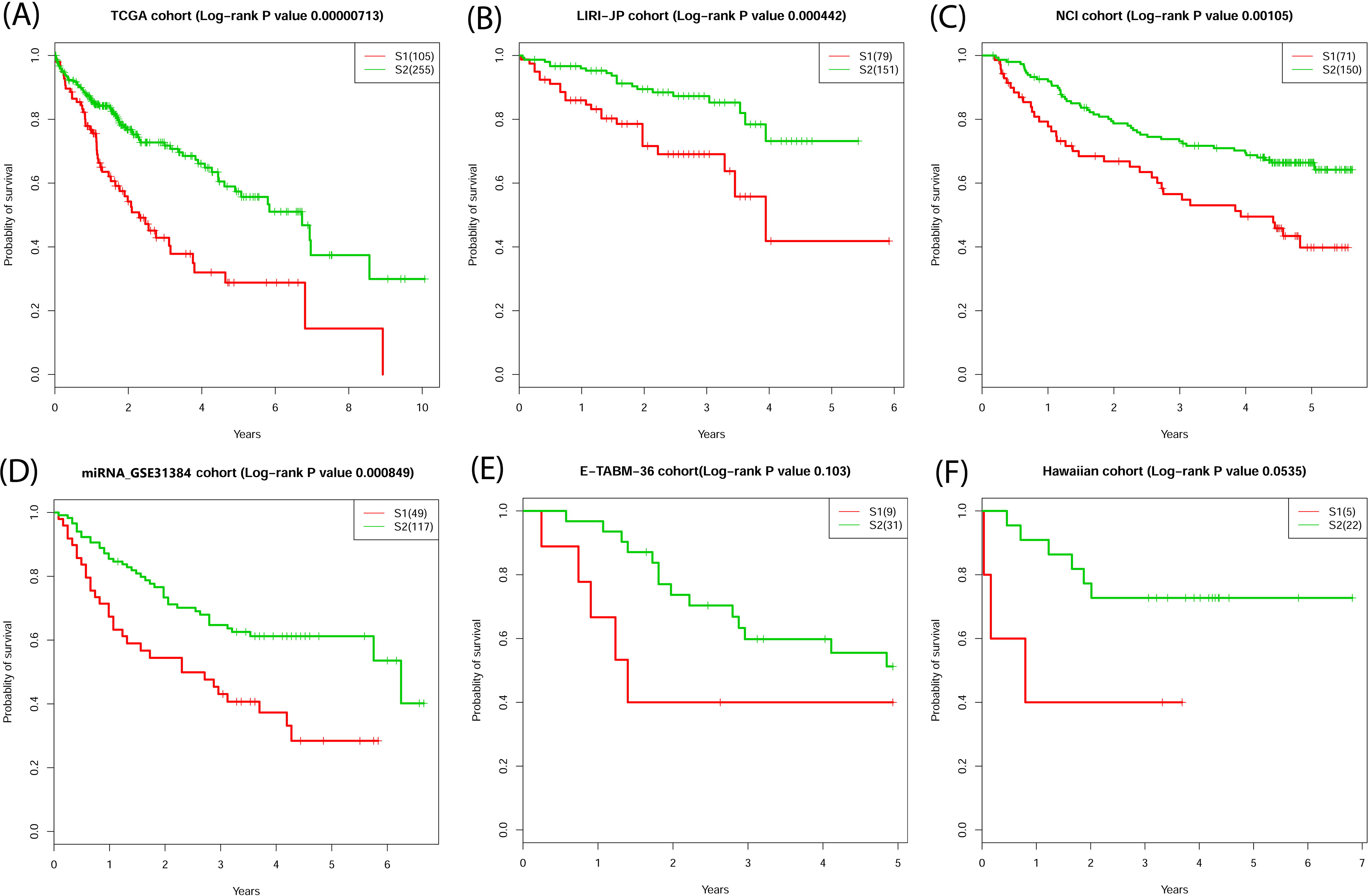
Significant survival differences for TCGA and external confirmation cohorts (A) TCGA cohort, (B) LIRI-JP cohort, (C) NCI cohort, (D) Chinese cohort, (E) E-TABM-36 cohort, and (F) Hawaiian cohort.

We next used the 2 classes determined above as the labels to build a classification model using the support vector machine (SVM) algorithm with cross-validation (CV) (Figure 1B). We split the 360 TCGA samples into 10 folds using 60/40 ratio for training and test data. We chose 60/40 split, rather than a conventional 90/10 split, in order to have sufficient test samples for sensible log-rank p-values in the survival analysis (see ‘Materials and Methods’). Additionally, we assessed the accuracy of the survival subtype predictions using C-index, which measures the fraction of all pairs of individuals whose predicted survival times are ordered correctly (42). We also calculated the error of the model fitting on survival data using Brier score (50). On average, the training data generated high C-index (0.70±0.04), low brier score (0.19±0.01), and significant average log-rank p-value (0.001) on survival difference (Table 1). Similar trend was observed for the 3-omics held-out test data, with C-index=0.69±0.08, Brier score=0.20±0.02, and average survival p-value=0.005 (Table 1). When tested on each single omic layer of data, this multi-omics model also has decent performances, in terms of C-index, low Brier scores and log-rank p-values (Table 1). These results demonstrate that the classification model using cluster labels is robust to predict survival-specific clusters. Table S2 enlists the top K features for 3-omics selected by ANOVA for the SVM-based classification in full TCGA cohort.

**Table 1:**
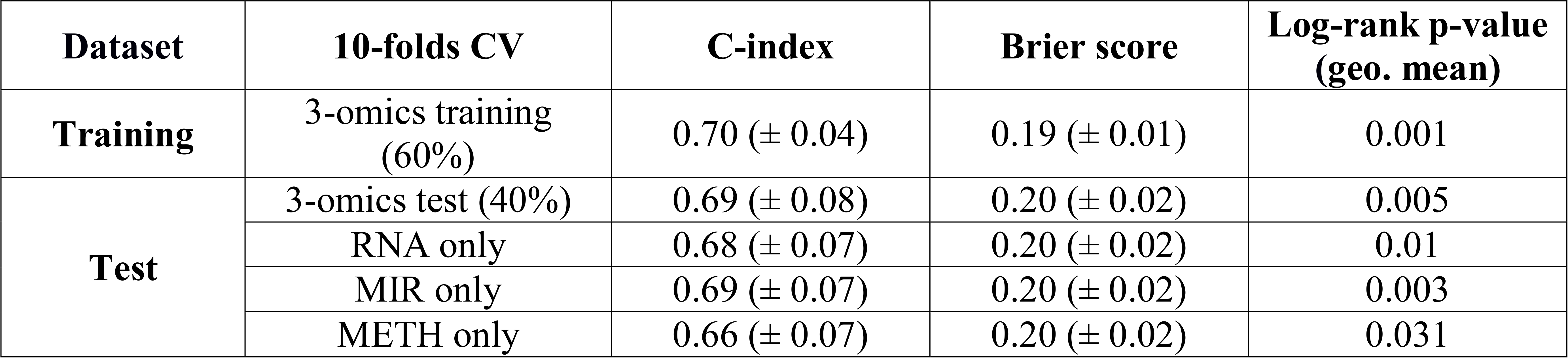
Cross-validation based performance robustness of SVM classifier on training and test set in TCGA cohort.

### The survival subtypes are robustly validated in five independent cohorts

To demonstrate the robustness of the classification model at predicting survival outcomes, we validated the model on a variety of five independent cohorts, each of which had only mRNA, or miRNA or methylation omics data (Table 2 and Figures 2B-2F). The common top features selected by ANOVA prior to SVM classification (between TCGA and 5 cohorts) are as follows: LIRI-JP (94%), NCI (74%), Chinese-GSE31384 (58%), E-TABM-36 (82%) and Hawaiian (100%). LIRI-JP dataset is the RNA-seq dataset with the most number of patients (n=230); we achieved a good C-index 0.75, a low Brier error rate of 0.16 and the log-rank p-value of 4.4e-4 between the two subtypes. For the second largest (n=221) NCI cohort (GSE14520), the two subgroups have decent C-index of 0.67 and low Brier error rate of 0.18 with log-rank p-value of 1.05e-3 (Table 2). For Chinese cohort (GSE31384), the miRNA array data with 166 samples, the two subgroups have C-index of 0.69, low Brier error rate of 0.21, and log-rank p-value of 8.49e-4 (Table 2). Impressively, the C-indices for the two smallest cohorts, E-TABM-36 (40 samples) and Hawaiian cohorts (27 samples) are very good, with values of 0.77 and 0.82, respectively. The p-values obtained for the small cohorts are not significant, due to small sample size, with values of 0.103 and 0.0535, respectively (Figures 2E, F).

**Table 2:**
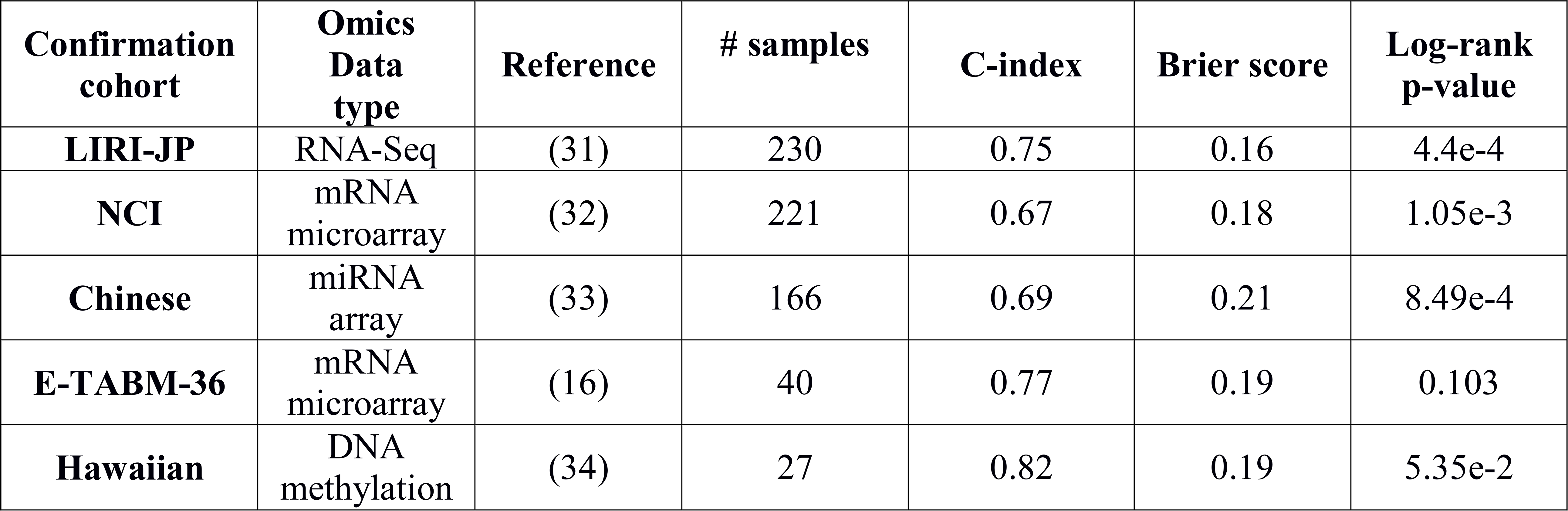
Performance of classifier for the five external confirmation cohorts.

### The DL-based methodology outperforms alternative approaches

We compared the performance of the model described in Figure 1B to two alternative approaches (Figure S2). In the first approach, we replaced autoencoder with the conventional dimension reduction method Principal Component Analysis (PCA). Similar to the 100 features from hidden nodes in the autoencoder, we obtained top 100 Principal Components, which were then subjective to univariate Cox-PH. As a result, 13 Principal component features remained. However, this approach failed to give significant log-rank p-value (P=0.14) in detecting survival subgroups (Figure S2A). It also yielded significantly lower C-index for test data (0.62) (Table S3), as compared to the model using autoencoder. In the second approach, we by passed the step of autoencoder, performed uni-variate Cox-PH analysis on each input feature in the 3 omics data types, and kept the top 37 features based on the C-index scores (Figure S2B). This model gave a P-value of 3.0e-8, still much less significant than the deep-learning method (6.0e-9, Figure S2C). More importantly, these alternative approaches failed overall to find significant subgroups in the majority of the confirmation sets. The only significance is in the LIRI-JP dataset using the Cox-PH approach (Table S3).

Worth noticing, the 3-omics based DL model gives better prediction metrics in CV, when compared to single-omics based DL models (Table S4), suggesting that indeed multi-omics data are better than single-omics data for model building. Finally, autoencoders fitted with a high number of epochs, with more than three hidden layers or with a high number of hidden nodes presented significant decreases of the performances. However, only one hidden layer or too few hidden nodes appeared also less efficient (Table S3).

### Adding clinical information does not improve DL-based multi-omics model

It remains to see if the DL-based multi-omics model will improve the predictability, by adding clinical information. Therefore, we assessed the performance of alternative models with clinical variables as the features, either alone or in combination with previous DL-based multi-omics model (Table 3). When clinical features were used as the sole feature set for survival prediction, the models’ performances were much poorer (Table 3), when compared to the DL-based genomic model (Table 2). Then we combined the clinical features with the 3 omics layers before the K-means clustering step in Figure 1B. Surprisingly, the C-indices of the combined model were not better on the confirmation cohorts with larger sample sizes (LIRI-JP and NCI cohorts), compared to those of DL-based multi-omics model. C-index and p-value were only slightly but not statistically significantly better for the Hawaiian cohort, which has only 27 samples. We thus conclude that the DL-based multi-omics model performs sufficiently well even without clinical features. We speculate the reason is due to the unique advantage of DL neural network, which can capture the redundant contributions of clinical features through their correlated genomic features.

**Table 3:**
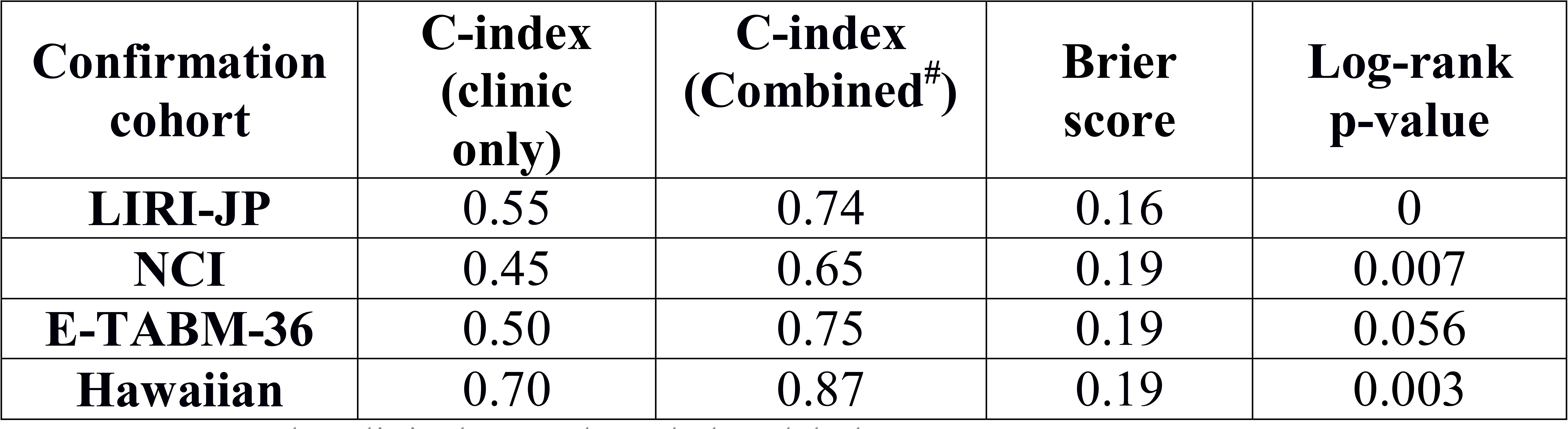
Performance of classifier for the five external confirmation cohorts.

### Associations of survival subgroups with clinical covariates

We performed the Fisher’s exact test between the two survival subgroups and the clinical variables from TCGA cohort, and found that only grade (P=0.0004) and stage (P=0.002) were significantly associated with survival, as expected. Since HCC is aggravated by the multiple risk factors including HBC, HCV, and alcohol, we also tested our model within subpopulations stratified by individual risk factors (Table 4). Impressively, our model performed very well on all the risk factor categories with C-indices ranging from 0.69-0.79, and Brier scores between 0.19 and 0.20. Log-rank P-values were significant in HBV infected patients (P=0.04), alcohol consumers (P=0.005) and others category (P=0.0035). The only non-significant p-value (P=0.20) was obtained from the HCV infected patients, probably attributed to the small group size (n=31).

**Table 4:**
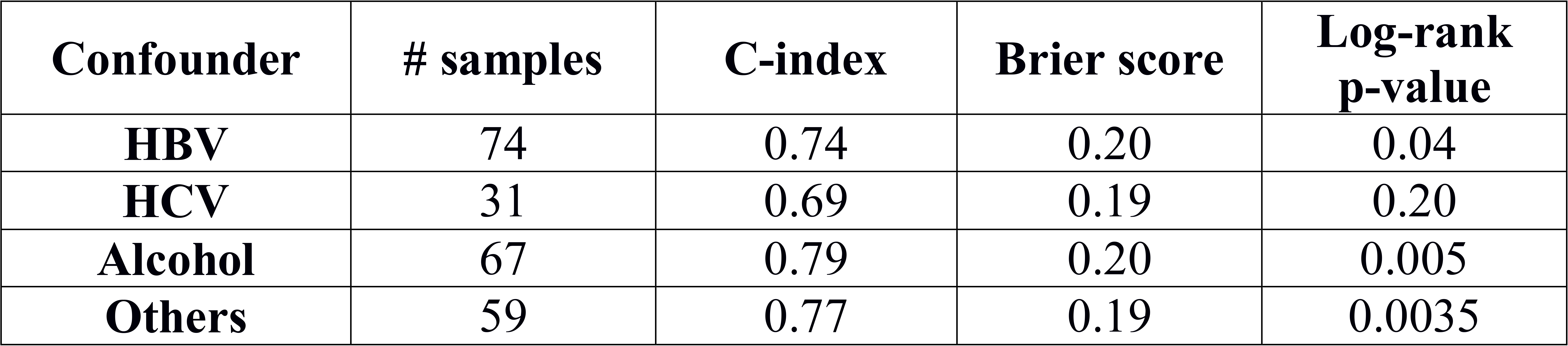
Performance of classifier for the five external confirmation cohorts.

*TP53* is one of the most frequently mutated genes in HCC, and its inactivation mutations have been reported to be associated with poor survival in the HCC (60). Between the 2 survival subgroups S1 and S2 in TCGA samples, *TP53* is more frequently mutated in the aggressive subtype S1 (Fisher’s test p-value=0.042). Further, *TP53* inactivation mutations are associated with the aggressive subtype S1 in LIRI-JP cohort, where whole genome sequencing data are available (p-value=0.024).

### Functional analysis of the survival subgroups in TCGA HCC samples

We used DESeq2 package (52) for differential gene expression between the two identified subtypes. After applying the filter of log2 fold change >1 and FDR <0.05, we obtained 820 up-regulated and 530 down-regulated genes in the aggressive sub-cluster S1. Figure 3 shows the comparative expression profile of these 1350 genes after normalization. The up-regulated genes in the S1 cluster include the stemness marker genes, *EPCAM* (P=5.7e-6), *KRT19* (P=6.7e-15) and tumor marker *BIRC5* (P=1.2e-13) genes, which were also reported earlier to be associated with aggressive HCC subtype (61-63). Additionally, 18 genes (*ADH1B*, *ALDOA*, *APOC3*, *CYP4F12*, *EPHX2*, *KHK*, *PFKFB3*, *PKLR*, *PLG*, *RGN*, *RGS2*, *RNASE4*, *SERPINC1*, *SLC22A7*, *SLC2A2*, *SPHK1*, *SULT2A1*, *TM4SF1*) differentially expressed in the two subtypes have similar trends of expression as in the previous study, where a panel of 65-gene signature was associated with the HCC survival (64).

**Figure 3:**
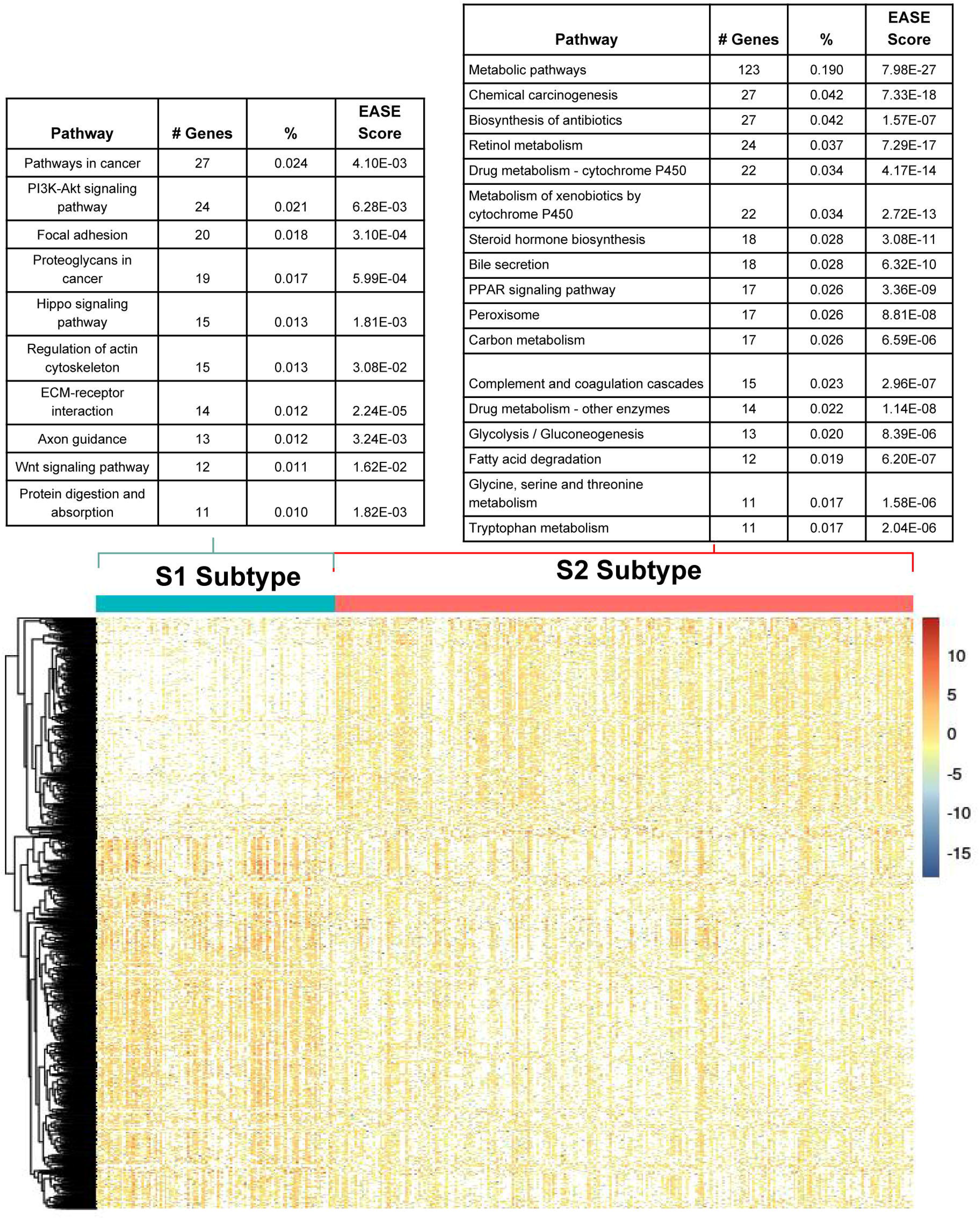
Differentially expressed genes and their enriched pathways in the two subtypes from TCGA cohort S1: aggressive (higher-risk survival) subtype; S2: moderate (lower-risk survival) subtype.

Using the differentially expressed genes above, we conducted KEGG pathway analysis to pinpoint the pathways enriched in two subtypes. Since we have used EASE score in DAVID as the enrichment method, these results should be interpretive only (65). These subtypes have different and (almost) disjoint active pathways, confirming that they are distinct subgroups at the pathway level (Figure 4). Aggressive subtype S1 is enriched with cancer related pathways, Wnt signaling pathway, PI3K-Akt signaling pathway etc. (Figure 4A). Wnt signaling pathway was reported being associated with aggressive HCC previously (66). In contrast, the moderate subtype S2 has activated metabolism related pathways including drug metabolism, amino acid and fatty acid metabolism etc. (Figure 4B). Further biological functional studies are needed to confirm the signaling pathway (for S1) vs. metabolic pathway (for S2) preferences between the two survival groups. We performed similar differential analysis for miRNA expression and methylation data, and detected 23 miRNAs and 55 genes’ methylation statistically different between the two subgroups (Figure S3 and File S2).

**Figure 4:**
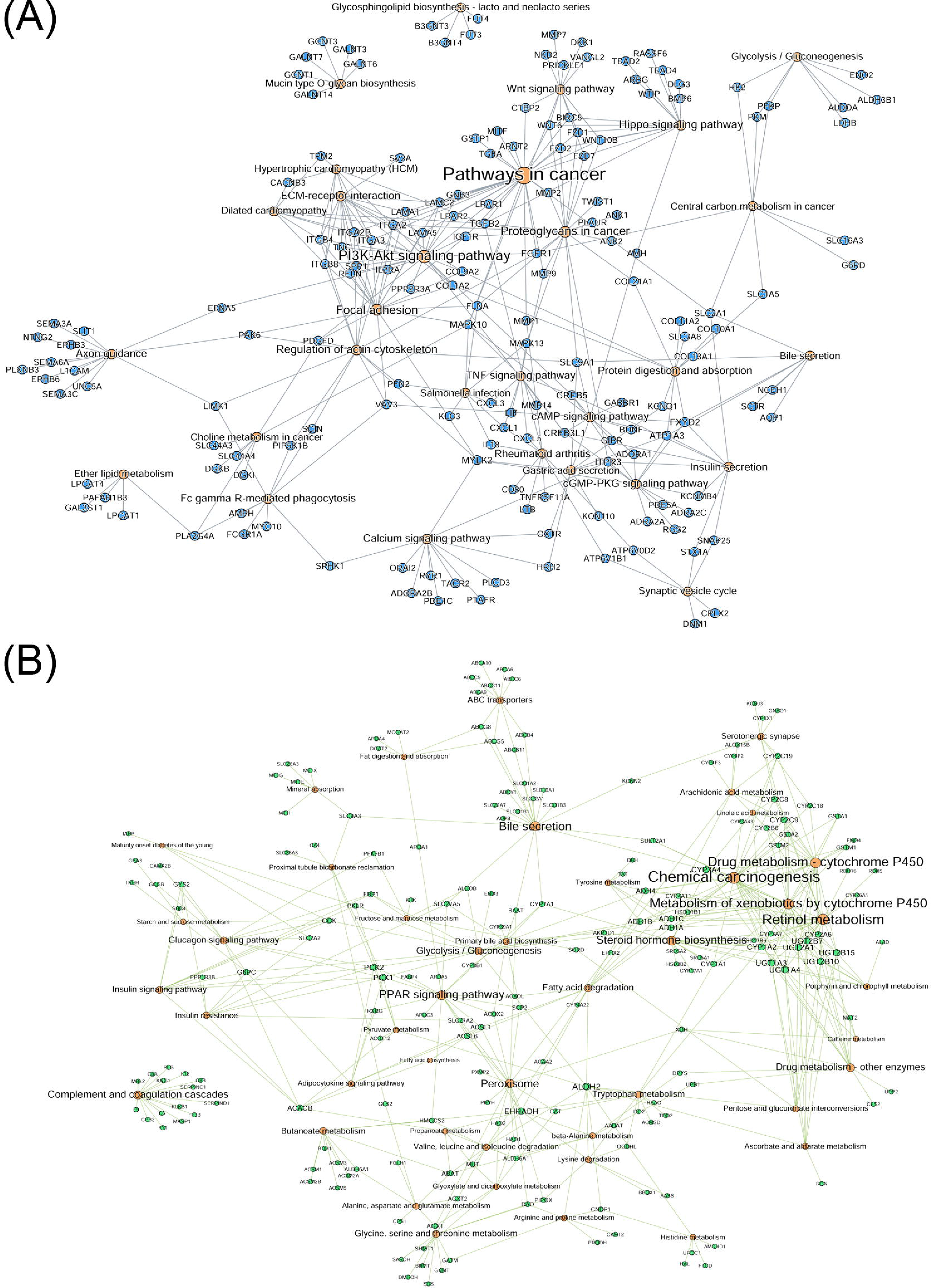
Bipartite graph for significantly enriched KEGG pathways and upregulated genes in two subtype. Enriched pathway-gene analysis for upregulated genes in the (A) S1 aggressive tumor sub-group and (B) less aggressive S2 sub-group.

## Discussion

Heterogeneity is one of the bottlenecks for understanding the HCC etiology. Though there are many studies for subtype identification of the HCC patients, embedding survival outcome of the patients as part of the procedure of identified subtypes has not been reported before. Moreover, most reported HCC subtype models have either no or very few external confirmation cohorts. This calls for better strategies, where the identified subtypes could reflect the phenotypic outcome of the patients i.e. the survival directly. Present work includes the integration of the multi-omics data from the same patients, giving an edge by exploiting the improved signal-to-noise ratio. To our knowledge, we are the first to use the deep learning framework to integrate multi-omics information in HCC. It propels deep learning to develop risk stratification model, not only for prognostication but also instrumental for improvising risk-adapted therapy in HCC.

We have identified two subtypes from the molecular level. This model is robust and perhaps more superior than other approaches, manifested in several levels. Firstly, CV results gave the consistent performance in TCGA HCC test samples, implying the reliability and robustness of the model. Secondly, deep-learning technique used in the model has captured sufficient variations due to potential clinical risk factors, such that it performs as accurately or even better than, having additional clinical features in the model. Thirdly, autoencoder framework shows much more efficiency to identify features linked to survival, compared to PCA or to individual Cox-PH based models. Lastly and most importantly, this model is repetitively validated in five additional cohorts, ranging from RNA-seq, mRNA microarray, miRNA array, and DNA methylation platforms.

In association with clinical characteristics, the more aggressive subtype (S1) has consistent trends of association with higher *TP53* inactivation mutation frequencies in the TCGA and LIRI-JP cohorts, which is in concordance with the previous study (60). Association of stemness markers (*KRT19*, *EPCAM*) with S1 subtype is also in congruence with the literature (61,62). Moreover, S1 subtype is enriched with activated Wnt signaling pathway (66). Despite our effort, the one to one comparison with the previous studies is not feasible due to the absence of cluster label information in original reports, and lack of survival data in some cases. Fortunately, we were able to identify five external confirmation cohorts encompassing different omic dataset, and succeeded in validating the subtypes among them. These results gave enough confidence that the 2 survival subtype specific model proposed in this report is of direct clinical importance, and maybe useful to improve HCC patients survival.

Some caveats are worth discussion below. First, we used whole TCGA data set in step 1 (Figure 1B), in order to learn the class label of the TCGA samples in an unsupervised way. Therefore, when we build a SVM model using TCGA training dataset and apply it on TCGA testing data, the C-statistics may be inflated; however, when we apply the SVM model to the other external datasets, these data sets give more unbiased C-statistics, since they are not part of the SVM model construction process. Also, our current model is trained on TCGA HCC data, and it has been reported earlier that TCGA samples are impure (67). Liver tumor samples (LIHC) was reported to have better than average purity among 21 tumor types, higher than Breast Cancers in TCGA. Also, to obtain only HCC samples, we have procured the data from the TCGA website with their clinical annotation for liver hepatocellular carcinoma under the LIHC flag. The purity issue, along with the heterogeneous nature of HCC due to various risk factor, may explain why we do not have C-index better than 0.80 in the TCGA training data. To further examine the effects of risk factors on the model, we built sub-models for samples with only HBV, HCV and alcohol risk factor. We obtained C-indices of 0.90, 0.92 and 0.83 on HBV, HCV and alcohol affected TCGA subpopulations. Thus the heterogeneity of the population does affect the model performance. However, issues exist to test these models on external datasets, since the submodels were built on small training data, thus they could suffer from over-fitting in confirmation cohorts. Moreover, sample risk-factors are not always known for public cohorts, restricting our confirmation effort. Albeit these issues, the current TCGA based model has an average C-index 0.74 on 5 external confirmation, indicating the model is generally predictive. Additionally, we used log-rank p-value and brier score as other performance metrics to assess our pipeline. In the future, we plan to collaborate with clinicians to prospective cohorts, and improve the model over time.

## Acknowledgements

This research was supported by grants K01ES025434 awarded by NIEHS through funds provided by the trans-NIH Big Data to Knowledge (BD2K) initiative (http://datascience.nih.gov/bd2k), P20 COBRE GM103457 awarded by NIH/NIGMS, NICHD R01 HD084633 and NLM R01LM012373 and Hawaii Community Foundation Medical Research Grant 14ADVC-64566 to LX Garmire.

## Authors Contributions

LXG envisioned the project, LL and KC prepared the dataset. OP, KC and LL developed the pipeline. KC, OP, LL and LXG wrote the manuscript. All authors have read, revised, and approved the final manuscript.

## Conflict of interest

The authors declared no conflict of interest.

